# Importance of linear combination modeling for quantification of Glutathione and GABA levels using Hadamard-edited MRS

**DOI:** 10.1101/2021.12.28.474256

**Authors:** Yulu Song, Helge J. Zöllner, Steve C.N. Hui, Georg Oeltzschner, James J. Prisciandaro, Richard A.E. Edden

## Abstract

**Purpose:** Two main approaches are used for spectral analysis of edited data: simple peak fitting and linear combination modeling (LCM) with a simulated basis set. Recent consensus recommended LCM as the method of choice for the spectral analysis of edited data. The aim of this study is to compare the performance of simple peak fitting and LCM in a test-retest dataset, hypothesizing that the more sophisticated LCM approach will improve quantification of HERMES data compared with simple peak fitting.

**Methods:** A test-retest dataset was re-analyzed using Gannet (simple peak fitting) and Osprey (LCM). These data were obtained from the dorsal anterior cingulate cortex of twelve healthy volunteers, with TE 80 ms for HERMES and TE 120 ms for MEGA-PRESS of glutathione (GSH). Within-subject coefficients of variance (CVs) were calculated to quantify between-scan reproducibility of each metabolite estimate.

**Results:** The reproducibility of HERMES GSH estimates was substantially improved using LCM compared to simple peak fitting, from a CV of 19.0% to 9.9%. For MEGA-PRESS data, the GSH reproducibility was similar using LCM and simple peak fitting, with CVs of 7.3% and 8.8% respectively.

**Conclusion:** Linear combination modeling with simulated basis functions substantially improves the reproducibility of GSH quantification for HERMES data.

## 1. Introduction

*J*-difference-edited ^1^H-MRS is a commonly used approach to quantify levels of low-concentration metabolites in the human brain^1^, including but not limited to γ-aminobutyric acid (GABA)^2–4^; N-acetyl aspartyl glutamate (NAAG)^5^; glutathione (GSH)^6^; ascorbate (Asc)^7^; aspartate (Asp)^8,9^; phosphorylethanolamine (PE)^10^ and lactate (Lac)^11^. Metabolites that are amenable to detection by *J*-difference editing have coupled spin systems and MR signals that are overlapped in the in-vivo spectrum. *J*-difference editing relies upon selective manipulation of the *J*-evolution of the spin system of the target metabolite. The efficiency of editing is therefore strongly dependent on the echo time (TE), with different spin systems having different optimal TEs for editing, e.g. GSH at TE ∼ 120 ms^12,13^ and GABA at TE ∼ 68 ms^2^. Recently, the MEscher–GArwood Point RESolved Spectroscopy (MEGA-PRESS) pulse sequence, which is limited to targeting one editing frequency per experiment, has been developed to Hadamard Encoding and Reconstruction of MEGA-Edited Spectroscopy (HERMES)^14^ for the selective detection of multiple metabolites.

The two most commonly edited metabolites are the major antioxidants glutathione GSH, and the primary inhibitory neurotransmitter GABA. As an antioxidant, glutathione plays a crucial role in protecting cells from oxidative damage by neutralizing reactive oxygen species – oxidative stress is an important mechanism in a range of neurodegenerative processes^15^. In-vivo detection of GABA is important for unraveling the role of inhibitory neurotransmission in healthy cortical processing^16^ and understanding the role of inhibitory dysfunction in a range of psychiatric^17^, neurological and neurodevelopmental disorders^18–20^. Thus simultaneous detection of GSH and GABA in a single HERMES experiment is a promising approach for efficiently measuring two potential biomarkers of key disease processes.

MRS is quantitative, in the sense that the size of signals detected is proportional to the number of spins and the concentration of molecules from which the signal derives. However, MR spectra require modeling to determine the size of signals, and in the common case of spectral overlap, to assign signal to particular metabolites. The two most commonly used approaches for modeling spectra are simple peak fitting and linear combination modeling (LCM). Simple peak fitting can be used to estimate an individual metabolite peak of interest using a simple, e.g. Gaussian, lineshape model. This method is commonly used for edited spectra which are usually relatively sparse. LCM uses a basis set to model different metabolite contributions to the full spectrum^21^, maximizing the prior knowledge leveraged while seeking to estimate a larger number of parameters. Recent consensus^21^ recommended LCM for quantification of edited MRS data, but there is relatively little literature to-date that supports that conclusion or identifies best practices.

The use of simple spectral modeling in the Gannet software^22^ is well-established for the quantification of GABA+-edited spectra (i.e. those in which there is a substantial macromolecular contribution mixed into the edited GABA signal)^23,24^. Although an analogous simple modeling approach has been developed for GSH, the properties of the spectrum to be modeled vary substantially with TE because of the strong TE modulation of adjacent co-edited aspartyl signals, and it was previously reported^25^ that modeling of HERMES data acquired at TE = 80 ms was less reproducible than for MEGA-PRESS acquired at TE = 120 ms. The aim of this study is to investigate whether this difference is indicative of a fundamental issue with multiplexed editing in HERMES, with GSH editing at TE = 80 ms, or with the modeling approach employed. We therefore compare the performance of simple peak fitting and LCM in this test-retest dataset, hypothesizing that the more sophisticated modeling approach will improve quantification of HERMES data compared with simple peak fitting.

## 2. Methods

### 2.1 MRS Acquisition

A test-retest reproducibility set of HERMES and MEGA-PRESS data has previously been acquired^25^. Data from twelve healthy volunteers measured in the dorsal anterior cingulate cortex (dACC) as shown in Figure 1 were acquired with TE = 80 ms for HERMES and TE = 120 ms for MEGA-PRESS of GSH. For simplicity, these acquisitions are referred to as MEGA-120 and HERMES-80 elsewhere in the manuscript. Acquisition methods are specified fully in the original manuscript^25^, and will only be briefly outlined here. Editing pulses (duration 20 ms) were applied to GABA spins at 1.9 ppm (for HERMES) and GSH spins at 4.56 ppm (for HERMES and MEGA-PRESS). Salient acquisition parameters were: TR = 2000 ms; 256 transients; 16-step phase cycling; spectral width 2.5 kHz; 2048 complex data points; 30 × 25 × 25 mm^3^ voxel in dACC, VAPOR water suppression^26^, internal water referencing^27^. HERMES editing schemes for the detection of GABA co-edit homocarnosine and macromolecular signals^14,28^; therefore, the edited 3-ppm GABA signal in the GABA-edited HERMES difference spectrum is denoted GABA+. A matched acquisition without water suppression was acquired for eddy-current correction and quantification^27^.

**Figure 1.**
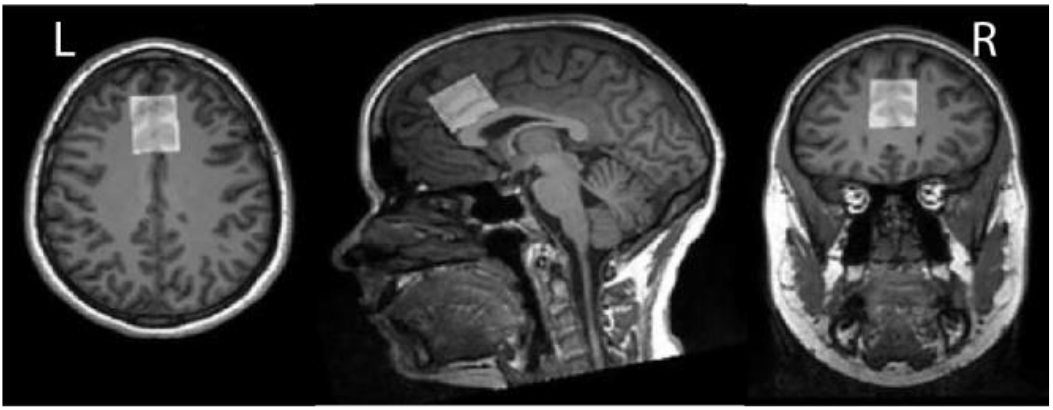
In vivo HERMES GSH- and GABA+-edited spectra were acquired from dorsal anterior cingulate cortex (dACC) voxel (30 × 25 × 25 mm^3^).

### 2.2 MRS Data Analysis

Simple peak fitting of the data was performed in Gannet (v3.1.5)^22^ (https://github.com/richardedden/Gannet3.1/releases/tag/v3.1.5) and LCM was performed in Osprey (v1.0.1.1)^29^ (https://github.com/schorschinho/osprey/releases/tag/v1.0.1.1), an open-source MRS analysis toolbox. Details about each implementation are expounded below.

#### 2.2.1 Gannet

Data processing in Gannet included spectral registration using a robust spectral registration approach for frequency-and-phase correction of the individual transients^30^, weighted averaging of the transients^30^, zero-filling to 32K data points, and 3-Hz exponential line broadening. The Gannet model for GABA+-edited spectra applies three Gaussian peaks to model GABA+ and the Glx doublet between 2.79 and 4.1 ppm, and a 4-parameter curved baseline function containing linear and quadratic terms. A five-Gaussian model is used for GSH-edited spectra acquired at TE = 80 ms, as for the HERMES data, and a six-Gaussian model for the GSH-edited spectra acquired at TE = 120 ms. In both cases, a 4-parameter curved baseline function containing linear and quadratic terms data is included, data are modeled between 2.25 and 3.5 ppm, and one Gaussian is assigned to model the GSH signal and the remainder to model the complex co-edited aspartyl multiplet at ∼2.6 ppm. Water reference data were quantified with a Gaussian-Lorentzian lineshape model. This Gannet analysis reproduces the analysis performed previously^25^.

#### 2.2.2 Osprey

The raw data were eddy-current corrected^31^ based on the water reference, and the individual transients were aligned separately within each sub-spectrum set (edit-ON or edit-OFF for MEGA-PRESS and sub-spectrum A, B, C, or D for HERMES) using robust spectral registration^30^. The averaged GSH MEGA-PRESS edit-ON and edit-OFF spectra were aligned by optimizing the relative frequency and phase such that the tNAA signal at 2 ppm in the difference spectrum was minimized. The HERMES sub-spectra were aligned in three pairwise steps, adjusting the frequency and phase such that the signal in different target regions is minimized in the difference spectrum: the residual water is minimized for the GSH-OFF sub-spectra (aligning B and D), the 2-ppm tNAA signal is subsequently minimized to align the GSH-ON-GABA-OFF sub-spectrum C to D, and finally the 3.2-ppm tCho is minimized to align sub-spectrum A to C. The final GSH-edited difference spectrum is generated by subtraction (MEGA-PRESS) or Hadamard combination (HERMES). A Hankel singular value decomposition (HSVD) filter^32^ was applied to remove residual water signals and to reduce baseline roll.

The basis sets were generated from a localized 2D density-matrix simulation (101 × 101 spatial grid, voxel size 30 mm × 30 mm, field of view 45 mm × 45 mm) implemented in a MATLAB-based toolbox FID-A^33^, using vendor-specific refocusing pulse shape and duration, sequence timings, and phase cycling. Nineteen metabolite basis functions were included (ascorbate, aspartate, creatine, negative creatine methylene, GABA, glycerophosphocholine, GSH, Gln, Glu, water, myo-inositol, lactate, NAA, NAAG, phosphocholine, phosphocreatine, phosphoethanolamine (PE), scyllo-inositol, and taurine). Spectra were modeled between 0.5 and 4.2 ppm. For the GSH-edited difference spectra, co-edited macromolecule (MM) peaks were parametrized as Gaussian basis functions at 1.2 and 1.4 ppm for MEGA-PRESS and HERMES. Similarly, two MM basis functions were added at 0.93 ppm and 3 ppm for the GABA+-edited HERMES difference spectrum. Values of GABA+ that are reported combine the signals modeled by the GABA basis function and the MM_3.0_ function. Amplitude ratio soft constraints are imposed on the MM and lipid amplitudes, as well as selected pairs of metabolite amplitudes, as defined in the LCModel manual^34^. Osprey’s default baseline knot spacing of 0.4 ppm was used for the spline baseline.

The water-reference data were quantified with a simulated water basis function in the frequency domain with a 6-parameter model (amplitude, zero-and first-order phase, Gaussian and Lorentzian line broadening, and frequency shift).

#### 2.2.3 Quantification

Metabolite estimates for GSH and GABA+ were quantified with respect to the unsuppressed water scan. No further relaxation or tissue-segmentation corrections were employed.

### 2.3 Statistical Analysis

The within-subject coefficient of variations (CVs) were calculated to quantify between-scan reliability of each metabolite^35^. Bland-Altman plots^36^ were used to visualize the agreement between metabolite levels from two consecutive scans.

## 3. Results

All data were successfully analyzed in Gannet and Osprey. The models in each case are presented in Figure 2B. It is worth noting the greater complexity of the LCM as well as the substantially different shape of the GSH model at 2.95 ppm and the different baseline handling. The variability in the models, visualized as the shaded region, is similar for Gannet and Osprey in quantifying MEGA data acquired at TE 120 ms. However, for HERMES data, LCM in Osprey appears to be less variable than Gannet with simple peak fitting. Metabolite levels quantified by the two algorithms for GSH and GABA+ are shown in Figure 3.

**Figure 2.**
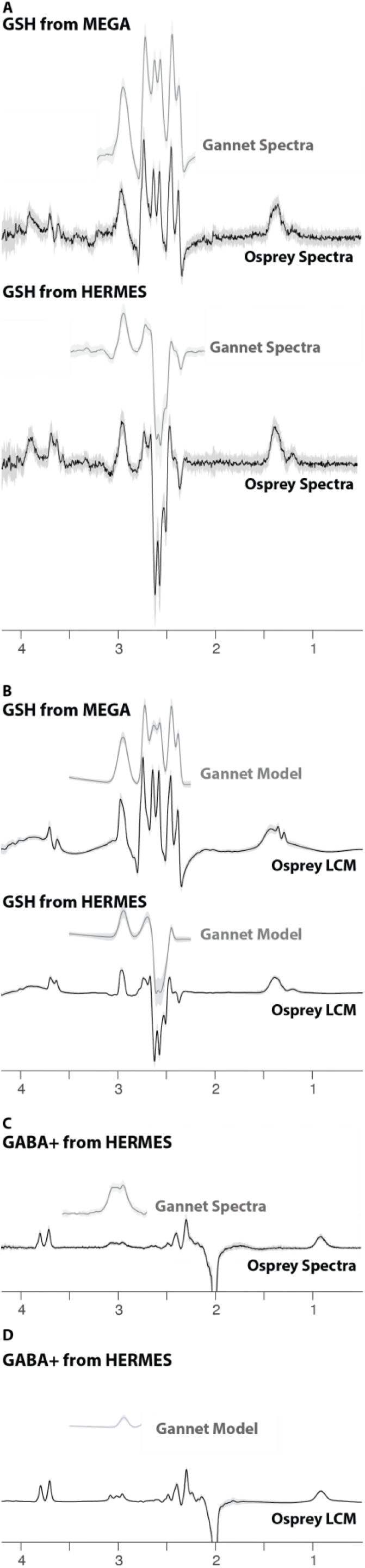
Edited difference spectra and models, using Gannet (gray) and Osprey (black) software. (A) Summary of GSH-edited data. (B) Summary of GSH models. (C) Summary of GABA+-edited data. (D) Summary of GABA+ models. The solid lines represent the group mean, woith the shaded area representing the range of mean ± one SD.

**Figure 3.**
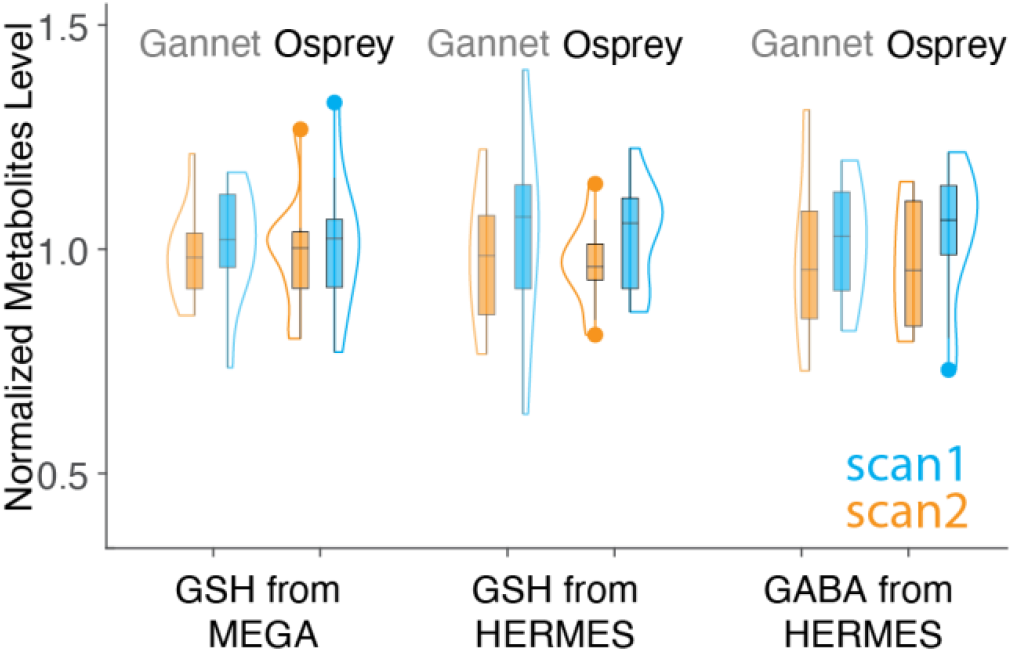
Violin plot of GSH and GABA+ estimates from simple peak fitting in Gannet and LCM in Osprey, grouped by sequence. Values for Gannet and Osprey have been normalized so that the mean of all values is 1, for the purposes of overlay. Note the greater variance of GSH_HERMES data using Gannet compared to Osprey.

Within-subject CVs are shown in Table 1. For GSH estimates, LCM showed similar levels of reproducibility for MEGA-PRESS and HERMES data (within-subject CVs of 9.9% and 8.8% respectively). Modeling of HERMES GSH spectra was more reproducible using LCM than with simple peak fitting (within-subject CVs of 9.9% and 19.0% respectively). For GABA+ estimates, the within-subject CV of the HERMES data using Osprey was 15.2%, which is comparable to previous estimates^23^ and simple peak fitting (16.7%).

**Table 1.** The within-subject CVs of HERMES data and MEGA-PRESS data modeled by Gannet and Osprey.

|  |  | **Gannet** | **Osprey** |
| --- | --- | --- | --- |
| GSH | MEGA-PRESS (TE = 120 ms) | 7.3%<sup>25</sup> | 8.8% |
|  | HERMES (TE = 80 ms) | 19.0%<sup>25</sup> | 9.9% |
| GABA+ | HERMES (TE = 80 ms) | 16.7%<sup>25</sup> | 15.2% |

Bland-Altman plots are shown in Figure 4, representing the agreement between scan 1 and scan 2 for subjects’ metabolite measurements. As shown in Panel A and C, the scatter dots were distributed within the 95% confidence interval. However, for the HERMES GSH data (Panel B), Osprey data were distributed within the 95% confidence interval, while Gannet data were distributed beyond 95% confidence interval, which indicates that modeling in Osprey improved the reproducibility compared to Gannet.

**Figure 4.**
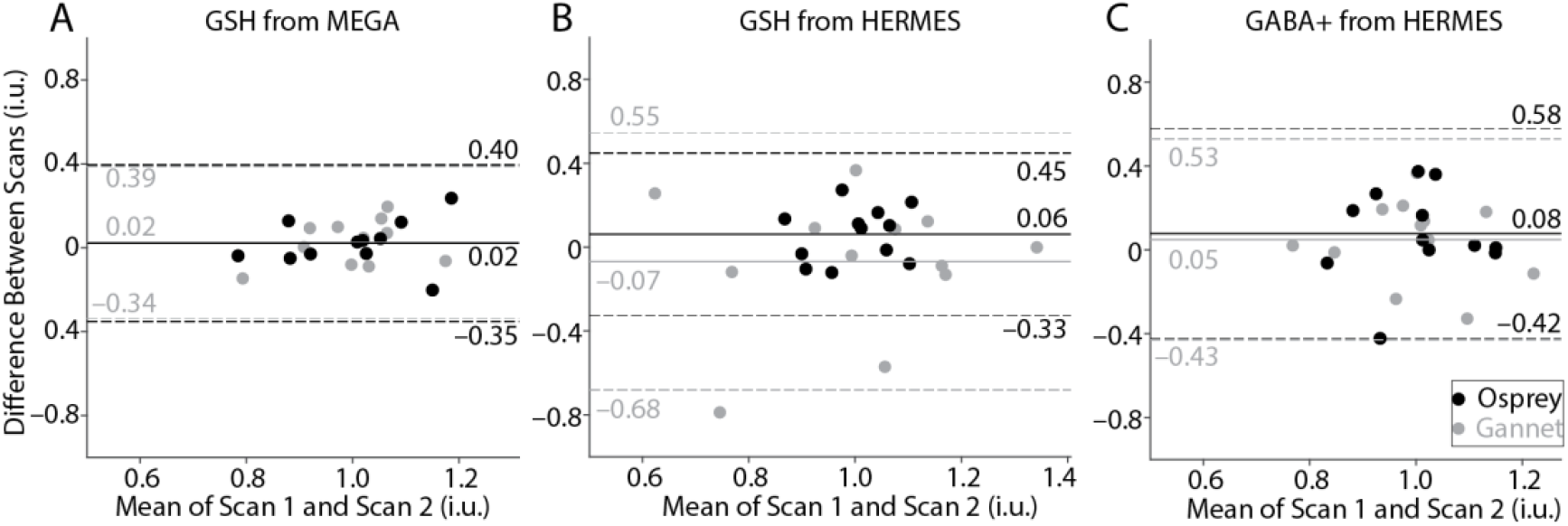
Bland-Altman plots of GSH-edited MEGA-PRESS (A) and HERMES (B) and GABA+-edited HERMES (C) data processed by Osprey (black) and Gannet (gray). Values for Gannet and Osprey have been normalized so that the mean of all values is 1, for the purposes of overlay. Solid lines represent the mean of the difference between scans, while dotted lines represent the 95% confidence interval.

## 4. Discussion

The reproducibility of quantification of GSH and GABA+ using linear combination modeling in Osprey was compared to simple peak fitting in Gannet. The main finding was that linear combination modeling with simulated basis functions substantially improved the reproducibility of GSH quantification with HERMES, compared to the simpler Gaussian modeling. The reproducibility of quantification of GABA+ using Osprey was in good agreement with Gannet. With improved modeling, the reproducibility of HERMES and MEGA-PRESS GSH measurements became similar, suggesting that HERMES is an appropriate choice for GSH quantification with the added benefit of performing GABA editing within the same experiment.

These findings suggest that the simple modeling approach employed in Gannet is not appropriate for GSH-edited spectra acquired at TE = 80 ms. In editing GSH, frequency-selective pulses are applied to the GSH signals at 4.56 ppm in order to resolve the GSH signal at 2.95 ppm in the difference spectrum. However, these pulses also invert aspartyl signals at ∼4.5 ppm, leading to co-editing of the coupled aspartyl signals at ∼2.6 ppm. These co-edited signals are complex multiplets (two doublets of doublets) that are wide and, particularly at shorter echo times (e.g., 80 ms), have complex mixed-phase lineshapes. While the Gannet model is sufficient for modeling one metabolite of interest using a simple Gaussian lineshape, it struggles to locate the appropriate baseline for these spectra. The Osprey model uses more prior knowledge as to the expected shape of the aspartyl signals, leading to greater control of the model baseline and more reproducible quantification of GSH.

In contrast, the aspartyl signals in MEGA-120 data are more consistently phased. The choice of TE in edited experiments depends on a number of factors: longer TEs can accommodate longer, more frequency-selective refocusing pulses^37^; shorter TEs suffer from less transverse relaxation; and the efficiency of editing is strongly dependent on the spin system of interest and the TE. MEGA-PRESS, in detecting one metabolite per experiment, offers greater flexibility in TE choice than HERMES, in which more than one metabolite is edited at the same TE value. In general, triplet-like signals such as GABA edit efficiently at medium TE ∼ 70 ms, whereas doublet-like signals such as GSH edit efficiently at long TE ∼ 140 ms. Importantly, when considering a TE-compromise for HERMES, triplet-like signals approach zero editing efficiency at these longer TEs. The optimal TE for editing GSH in phantoms is ∼120 ms, although owing to T_2_ relaxation, the in-vivo edited signal is not substantially different between TEs of 68 and 120 ms^13^. However, the edited spectra are substantially different, largely due to the differing behavior of the aspartyl signals. It has previously been noted^12^ that this leads to spectra that are more challenging to quantify, and indeed pioneering work applying linear combination modeling to edited spectra^6^ focused on GSH- and Asc-edited spectra. Medium-TE GSH editing offers signals that are less heavily T_2_-weighted, which is itself important for quantification, particularly in studies where relaxation rates may differ across comparison conditions (e.g., aging^38^). Linear combination modeling is more robust to complex signal lineshapes than simpler modeling.

This dataset has a number of inherent limitations. In comparing MEGA-120 and HERMES-80, two factors are changing (both MEGA/HERMES and the TE) making interpretation of the results more complex. Our interpretation of the data tends to focus on the TE difference, but it is also possible that there are inherent differences between HERMES and MEGA. Additionally, this study compared two widely used models of edited MRS spectra: simple peak fitting and linear combination model with simulated basis set. We could have included additional algorithms to increase the understanding of different modeling methods. However, those comparisons would be overwhelming and beyond the scope of a single study. Thirdly, it is notable that the within-subject CV of GABA was higher than seen on average for between-subject CVs in a recent multi-site GABA+ study^39^. This may reflect the limited estimation of variance in a single study with a low sample size (and these values are within the range of prior work), but also the different acquisition parameters. Compared to TE 68 ms, TE-80 GABA+ data have lower SNR due to slightly reduced editing efficiency and greater T_2_ relaxation of GABA and particularly MM signals. The longer editing pulses used also reduce the extent of MM co-editing, further impacting GABA+ SNR, and potentially introduce greater susceptibility to frequency instability. Any methodological changes that reduce the MM contamination to the GABA+ signal tend to negatively impact reproducibility.

## 5. Conclusion

When using linear combination modeling, GSH levels showed comparable reproducibility between HERMES-80 and MEGA-PRESS-120. This finding differs substantially from our previous analysis of this dataset with simple peak modeling. The advantages of HERMES, in terms of acquiring edited GSH and GABA+ spectra in a single acquisition, do not come at the cost of worsened GSH reproducibility when linear combination modeling is used in place of simple peak modeling. Linear combination modeling based on prior knowledge metabolite basis spectra is important for reproducible analysis of medium-TE GSH-edited spectra.

